# Evolution of novel sensory organs in fish with legs

**DOI:** 10.1101/2023.10.14.562285

**Authors:** Corey AH Allard, Amy L Herbert, Stephanie P Krueger, Qiaoyi Liang, Brittany L Walsh, Andrew L Rhyne, Allex N Gourlay, Agnese Seminara, Maude W Baldwin, David M Kingsley, Nicholas W Bellono

**Author notes:** Equal contribution (Corey AH Allard, Amy L Herbert, Stephanie P Krueger).

## Abstract

How do animals evolve new traits? Sea robins are unusual “walking” fishes that use leg-like appendages to navigate the seafloor. Here, we show that legs are *bona fide* sense organs that mediate the unique ability to localize and uncover buried prey. We then probe the developmental and physiological basis of these novel sense organs as a striking example of a major trait gain in evolution. We find certain sea robin species have legs with unique end-organs called papillae that mediate enhanced mechanical and chemical sensitivity to enable predatory digging behavior. Papillae exhibit dense innervation from touch-sensitive neurons, noncanonical epithelial taste receptors, and chemical sensitivity that drives predatory digging behavior. Using a combination of developmental analyses, crosses between species with and without papillae, and interspecies comparisons of sea robins from around the world, we demonstrate that papillae represent a key evolutionary innovation associated with behavioral niche expansion on the seafloor. These discoveries provide a conceptual framework for understanding how molecular, cellular, and tissue-scale adaptations integrate to produce novel organismic traits and behavior.

## Main

The evolution and diversification of novel traits enables adaptive behavior and expansion into new ecological niches. Yet, there are few examples that connect the molecular and developmental origins of a major trait gain with its physiological, behavioral, and ecological relevance. One remarkable example of trait gain is the “leg”-like appendage found within the family of fishes known as sea robins (Triglidae). Legs consist of six independently controlled, detached pectoral fin-rays used for “walking” and are thought to facilitate predation of food sources concealed beneath the seafloor^1–4^. Indeed, sea robins are so adept at finding prey that other fishes follow them to steal otherwise undetectable potential food sources^5^. By exploring this largely unstudied system, we made several serendipitous discoveries which define sea robin legs as novel sense organs that mediate species-specific functions suited to distinct behavioral niches.

## Sea robins use sensory legs for digging

Anecdotal stories of sea robins’ ability to find buried prey are common in the fishing community, but this behavior has not been examined experimentally. To test this ability, we developed a simple behavioral assay in which sea robins (*Prionotus carolinus*) were housed in a controlled tank with either mussels or capsules containing crude or filtered mussel extract buried in sand without visual cues (**Fig. 1a, b, Supplementary movie 1**). Sea robins alternated between short bouts of swimming and walking (**Fig. 1b**) and appeared to “scratch” at the sand surface with their legs while walking, which we hypothesized represented sensory behavior. Indeed, sea robins regularly found all buried prey-related items but did not uncover control capsules containing sea water (**Fig. 1c**), consistent with previous physiological evidence that the legs of some sea robins respond to chemical, tactile, and proprioceptive stimuli^2,6^. To determine if leg sensation facilitates predatory digging behavior, we tested whether the leg nervous system transduces specific cues that evoke sensory behavior. Using electrophysiological recordings from spinal nerves that innervate legs, we found that distal legs were robustly mechanosensitive (**Fig. 1d, e**). Furthermore, legs responded to prey extracts and common tastants for fish such as small non-polar L-amino acids^7^, and other amine and imine molecules including D-amino acids, methylated amino acids, and GABA-related compounds (**Fig. 1f, Extended Data Fig. 1a**). We also measured responses to compounds typically associated with nociception, such as TRP channel agonists, which elicited slower response kinetics compared with tastants (**Fig. 1f**). Based on this physiological profile, we repeated behavioral experiments using capsules containing single molecules that evoke leg neural activity. Remarkably, sea robins successfully located capsules containing only single leg agonists such as L-alanine, betaine, or D_2_-aminobutyric acid (D_2_ABA), but not TRP channel agonists or inert compounds (**Fig. 1g**). These results demonstrate that leg chemosensitivity for tastant-like molecules contributes to predatory behavior and suggest that legs could function as sensory organs.

**Figure 1.**
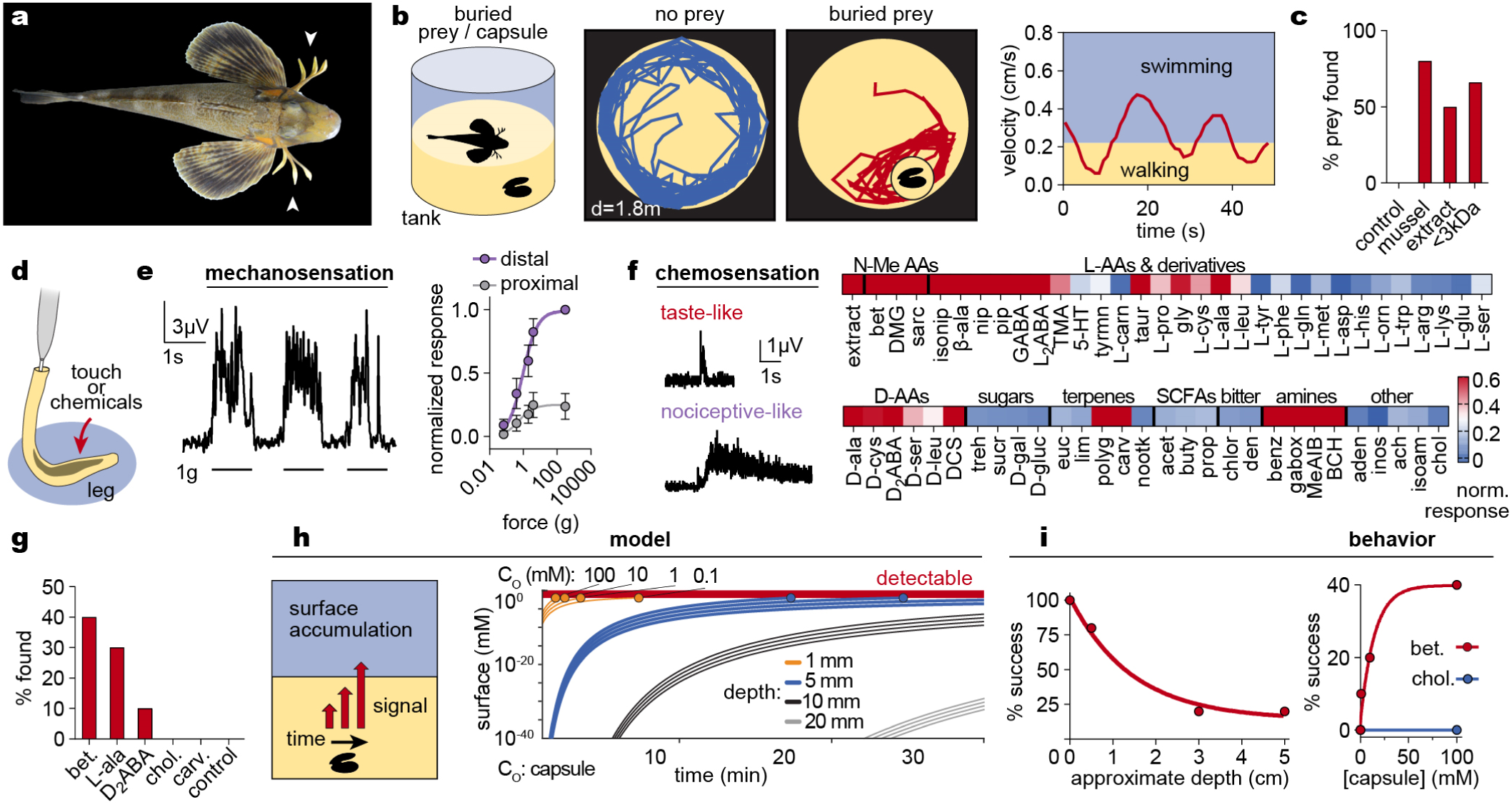
Sea robins are fish with legs that sense and find buried prey. **a,** Sea robins (*Prionotus carolinus*) are fish with leg-like appendages (arrowheads). **b,** Sea robins explored tanks with buried mussels by alternating between swimming (fast) and walking on sand (slow). Representative traces from 10 experiments. **c**, Sea robins localized and uncovered buried mussels, or capsules containing mussel extract or <3kDa filtered extract, but not capsules containing control sea water (*n* = 10 trials each). **d,** Schematic of electrophysiological recording from leg-specific spinal nerves. **e,** Distal legs responded to mechanical stimulation from von Frey filaments of varying stiffness. *n* = 4 legs with 4-10 stimulations per filament per leg, p < 0.0001 for comparison of curve plateau by sum-of-square F-test, plateau = 0.8342 to 1.163 (distal) vs. 0.1959 to 0.3132 (proximal). Data represented as the mean ± s.e.m. **f**, Legs responded to a broad range of chemicals including common tastants, marine osmolytes, and TRP channel agonists. Heatmap of relative responses from >6 legs. Responses to tastants (2 mM L-alanine) exhibited markedly faster kinetics compared to those elicited by TRP channel agonists (1 mM carvacrol). Recording representative of 6 recordings. Abbreviations: N-Me AAs: N-methylated amino acids, L-AAs: L-amino acids, D-AAs: D-amino acids, SCFAs: short chain fatty acids. **g**, Sea robins found capsules containing solutions of single leg agonists but not inert chemicals (*n* = 10 experiments per chemical). **h**, Model of chemical diffusion and surface availability in which shallow capsules of high concentrations produce detectable surface chemicals. **i**, Sea robins found mussels buried at shallow depths and higher concentrations. *n* = 10 trials each.

How do sea robins use their legs to sense prey if they are covered by sand? Chemicals released by prey would need to diffuse through sand and accumulate at sufficient surface concentration for sensation. To understand the limitations of detection, we developed a mathematical model that describes diffusion in sand over a spatial scale that depends on time and the physicochemical qualities of stimuli emanating from buried prey sources. Using values consistent with our behavioral experiments, we found that stimulus sources must be concentrated and shallow to reach the surface without being significantly diluted (**Fig. 1h**). To test this prediction, we asked if sea robins could find mussels buried at varied depths and concentrations. As expected, the success rate of capture sharply decreased with depth (**Fig. 1i**). Furthermore, sea robins were successful at finding capsules containing leg agonists ranging from 1 mM to 100 mM, corresponding with micromolar surface concentrations (**Fig. 1i**). Thus, sea robin legs detect buried chemical sources using near-surface cues at micromolar concentrations, consistent with their benthic ecology and leg chemosensitivity.

## Digging sea robins have specialized legs

During the course of behavioral experiments, we accidentally obtained a second species of sea robin, *Prionotus evolans,* which also has legs (**Fig. 2a**). Remarkably, we found that *P. evolans* failed to find buried mussels, squid, or crabs, despite consuming the same prey when visibly presented (**Fig. 2a**). Based on these observations, we conclude that *P. carolinus* is a robust digging species while *P. evolans* is non-digging and uses its legs for locomotion and probing visible prey. Considering the prominent role of sensory legs in digging behavior, we wondered whether interspecies behavioral differences reflect leg specialization. To our surprise, we observed striking macroscopic differences in leg morphology between the two species. The distal legs of digging *P. carolinus* were shovel-shaped and covered in epithelial protrusions similar to papillae found on tongues and along the gastrointestinal tract of many animals (**Fig. 2b**). In contrast, non-digging *P. evolans* legs were rod shaped, lacked papillae, and instead exhibited broad, shallow ridges in regions corresponding to papillae-covered surfaces in *P. carolinus* (**Fig. 2b**). Based on these morphological differences, we hypothesized that papillae are specializations that facilitate digging behavior.

**Figure 2.**
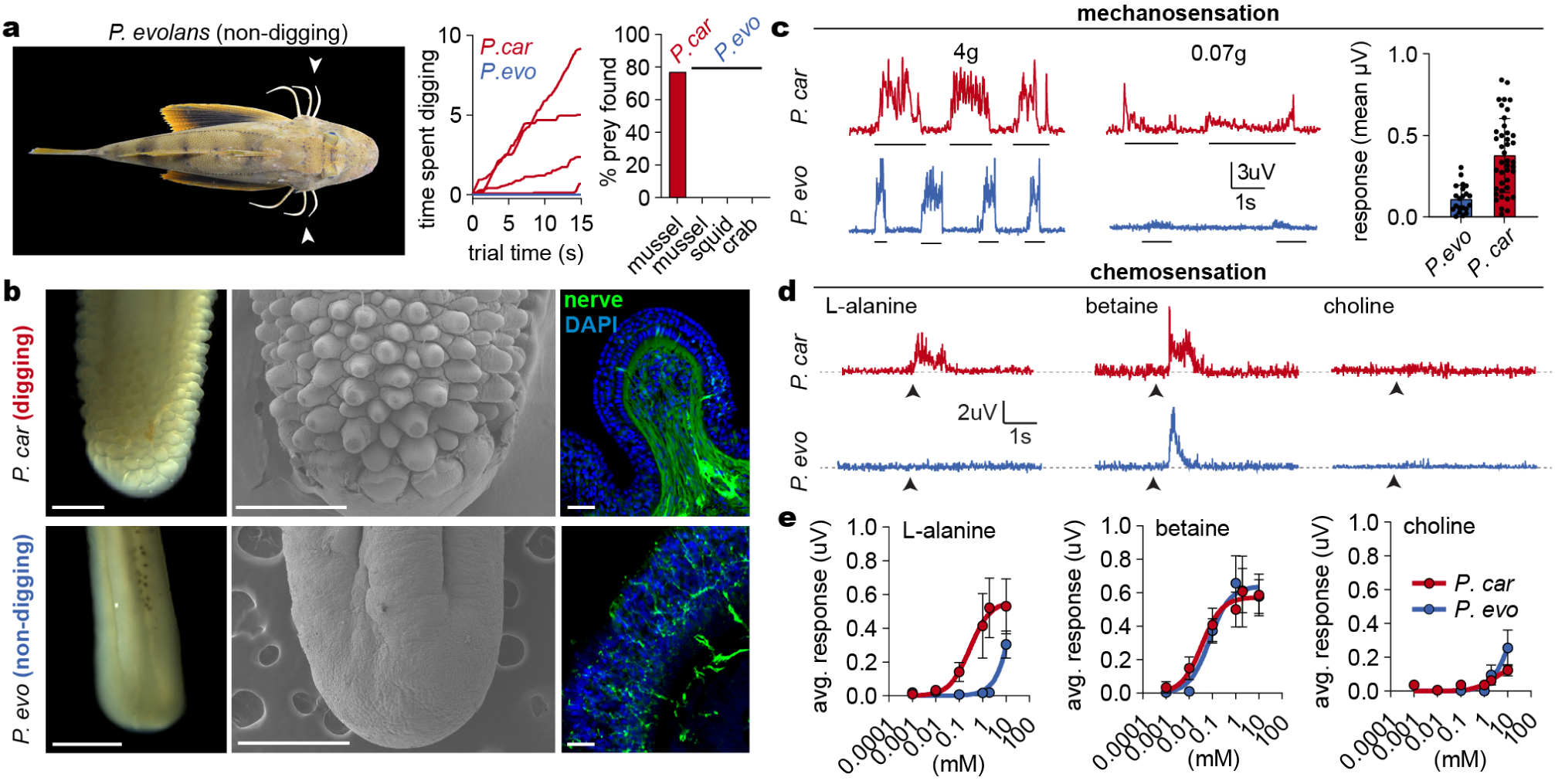
Legs of digging sea robins are specialized sensory organs. **a,** (*left*) *Prionotus evolans*. (*right*) *P. carolinus* but not *P. evolans* exhibited digging behavior and found buried prey (*n* = 10 mussel trials per indicated prey per species, 4 representative traces of mussel experiments for both species). **b**, Distal legs of digging *P. carolinus* were covered in papillae, which were absent in non-digging *P. evolans*. (*Left*) brightfield, (*middle*) scanning electron microscopy, (*right*) immunofluorescence for neural antigen HNK-1 (green) revealed that leg papillae and ridges are densely innervated in *P. carolinus* and modestly in *P. evolans,* respectively. Scale bars (1 mm left, 500 µm middle, 25 µm right). **c**, Legs of both species responded to strong mechanical stimulation (4g filament), but only *P. carolinus* responded to light mechanical stimulation (0.07g filament, *n* = 41 *P. carolinus*, 22 *P. evolans*, *p* <0.0001, t-test with Welch’s Correction). **d**, Sensation of L-amino acids (2 mM) was unique to digging *P. carolinus*, while both species responded to the marine osmolyte betaine (2 mM) and showed little response to choline (2 mM). Representative nerve recordings of >5 legs. **e**, Digging *P. carolinus* legs are ∼100X more sensitive to L-amino acids than non-digging *P. evolans* (responses detected at 100 µm in *P. carolinus* vs 10 mM *P. evolans*). *n* = 4 legs. Data in **c** and **d** represented as the mean ± s.e.m.

Leg papillae resemble sensory structures such as taste bud-containing oral papillae and mechanosensory papillae found on the facial regions of animals ranging from birds to star-nosed moles^8^. However, leg papillae lacked markers for canonical chemosensory cell types such as those in taste buds, solitary chemosensory cells, and olfactory receptor neurons, which we could readily detect in other sea robin tissues (**Extended Data Fig. 1b-f**). Instead, the only putative sensory cells we observed within papillae were free nerve-endings that permeate the distal leg epithelium in both *P. carolinus* and *P. evolans*. Nerve endings were densely concentrated within individual papilla in *P. carolinus,* but distributed along the broad ridges of *P. evolans* legs^6^ (**Fig. 2b**). Consistent with established function of free nerve endings, the distal legs of both sea robin species were sensitive to strong mechanical stimuli. However, papillae-covered surfaces of *P. carolinus* exhibited enhanced touch sensitivity that could reflect an adaptation that allows digging species to differentiate among objects, textures, or motion in sand (**Fig. 2c, Extended Data Fig. 2**).

Chemical stimuli are required for sensory digging behavior, so we next wondered whether chemosensitivity is distinct in digging sea robins. Whereas digging *P. carolinus* exhibited robust responses to common appetitive chemicals like L-amino acids, non-digging *P. evolans* legs were insensitive and instead only responded to a subset of chemicals, including nociceptor-associated molecules (**Fig. 2d, Extended Data Fig. 1a**). Remarkably, dose-response experiments revealed that the legs of digging sea robins were ∼100X more sensitive to L-amino acids than non-digging species, consistent with behavioral results and limitations of stimulus diffusion (**Fig. 2e**). Thus, enhanced chemical sensitivity in *P. carolinus* legs is associated with the ability to localize buried prey.

## Molecular and cellular basis of sensory adaptation

What are the molecular and cellular mechanisms of enhanced chemosensation in digging sea robins? Free nerve endings are typically associated with mechanosensation and chemesthesis (thermal, nociceptive, and tactile sensations)^9^, but not the detection of appetitive stimuli like the amino acids that elicit digging behavior. Early retrograde tracing of nerve endings in legs demonstrated that neuron cell bodies reside in dramatically enlarged spinal dorsal root ganglia (DRG), which connect to six novel spinal cord lobes that are specific to each leg^10^ (**Fig. 3a**). This is consistent with leg nerve responses to sensory agonists including the common tastant L-alanine, and the Piezo mechanoreceptor agonist Yoda1, which induced responses with kinetics similar to those recorded from nociceptor agonists (**Fig. 3b**, **Fig. 1h**). To characterize cellular properties, we dissociated and cultured functional sensory neurons from the specialized ganglia of digging *P. carolinus* (**Extended Data Fig. 2**). Whereas isolated neurons responded to Yoda1, the nociceptive TRP channel agonist carvacrol, and depolarizing high-extracellular potassium, we did not record measurable responses to appetitive leg agonists such as L-alanine. (**Fig. 3c, d**). Furthermore, most neurons exhibited mechanical stimulus-induced currents that varied in kinetics, suggesting a diversity of mechanoreceptor subtypes which is indicative of specialized somatosensory function (**Fig. 3e**). Thus, we conclude that sensory ganglia neurons mediate leg responses to touch and nociceptor-associated molecules but not to appetitive chemicals.

**Figure 3.**
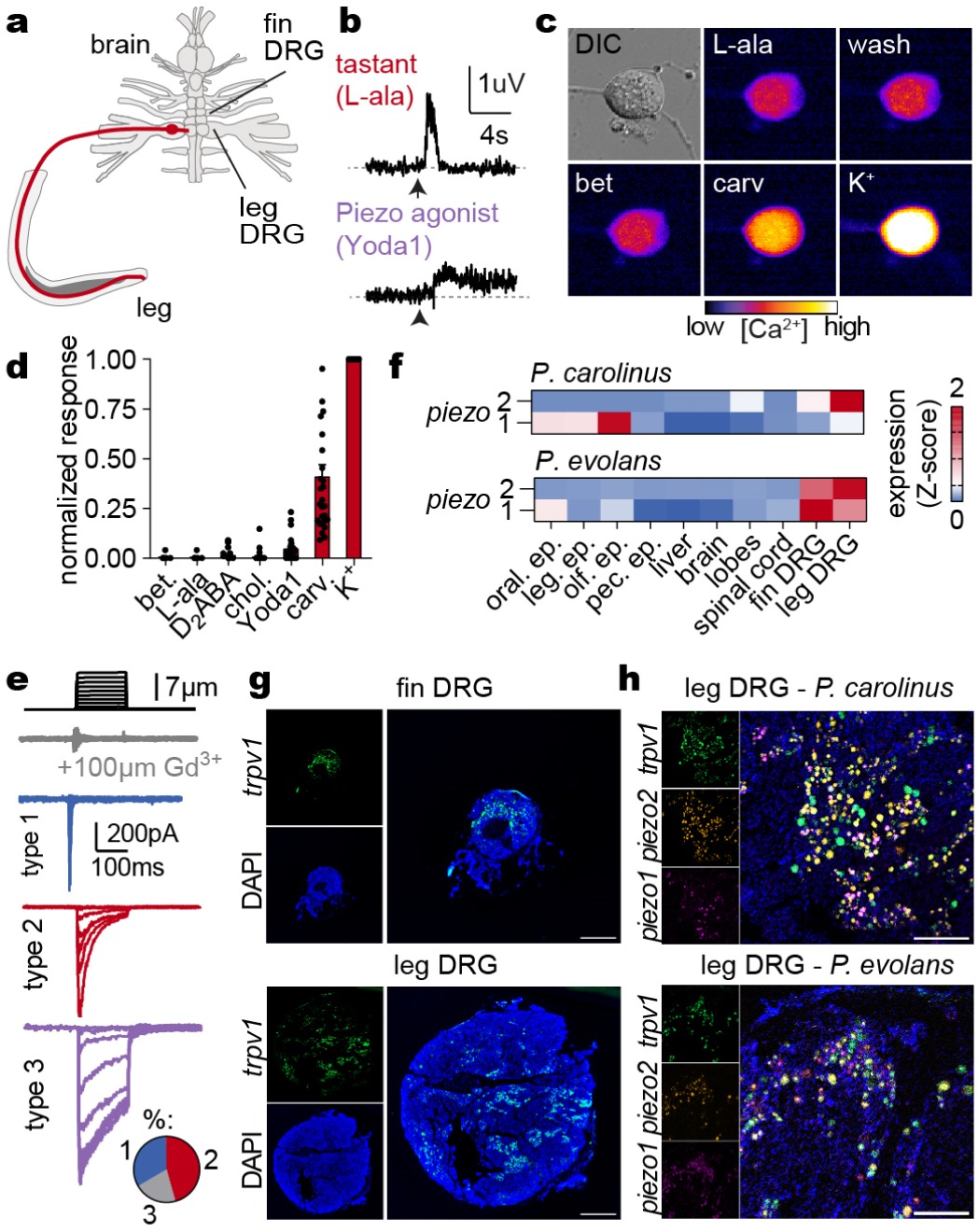
Sensory neurons mediate leg mechanosensation. **a,** Diagram of sea robin leg neural architecture^10^. **b**, Responses to 2 mM L-alanine had faster kinetics compared to those elicited by the Piezo mechanoreceptor agonist Yoda1 (5 µM). Representative of 6 recordings. **c-d**, Cultured sensory neurons from *P. carolinus* leg ganglia responded to depolarizing K^+^ (70 mM), the TRP channel agonist carvacrol (100 µM), and weakly to the Piezo agonist Yoda1 (5 µM), but not to appetitive (10 mM betaine, L-alanine, L-proline, D_2_ABA) or control chemicals (choline 10 mM) (*n* = 28 cells, one-way ANOVA *p* < 0.0001 carv. and K^+^). Data represented as the mean ± s.e.m. **e**, Sensory neurons were mechanosensitive and could be separated into three functional populations (*n* = 26, 8 type 1 with fast desensitization, 11 type 2 with intermediate desensitization, and 5 type 3 with slow desensitization). **f**, *piezo* mechanoreceptor mRNA transcripts were enriched in leg ganglia of both species. Scale: z-scaled normalized counts. Abbreviations: oral ep: oral epithelium, leg ep: leg epithelium, olf. ep: olfactory epithelium, pec. ep: pectoral fin epithelium. **g-h**, Leg-specific spinal ganglia were enlarged relative to fin ganglia and possessed an expanded population of *trpv1*-positive sensory neurons (green, DAPI in blue, scale bar 500 µm). Large populations of ganglia sensory neurons expressed mechanosensitive *piezo1* (magenta) and *piezo2* (orange) ion channels (DAPI in blue, scale bar = 200 µm). Images representative of 3 animals per species.

We next analyzed the molecular basis of sensory leg function. Consistent with mechanosensitivity of legs and the sensory neurons that innervate them, leg ganglia from both species exhibited enriched expression of the mechanosensitive ion channels *piezo1* and *piezo2* but did not express chemoreceptors for appetitive molecules (**Fig. 3f**). Both *piezo* channels colocalized with the sensory neuron marker *trpv1* (**Fig. 3g, h**) and *piezo1* transcripts were particularly enriched in neurons that innervate legs compared with those from pectoral fins (**Fig. 3g, h, Extended Data Fig. 2**), suggesting that *piezo1* may serve an unappreciated role in environmental sensation in fishes^11,12^. Considering that the leg-specific, enlarged dorsal root ganglia of both species contain sensory neurons expressing similar mechanosensitive ion channels, we conclude that mechanosensation across species shares a common molecular and cellular basis and suggest tissue-level adaptions (papillae versus ridges) underly species-specific mechanosensory properties.

If sensory neurons are not responsible for appetitive chemosensation, we wondered if this modality could be mediated by noncanonical sensory cells within the leg epithelium. To determine whether specific leg regions are specialized for chemosensation, we used a split-recording chamber and found that only the papillae-covered distal leg was responsive to appetitive chemicals (**Fig. 4a**). We then exploited this anatomical specialization to identify candidate chemoreceptors using comparative transcriptomics from distal versus proximal legs. Strikingly, the taste receptor *t1r3* was the single most-upregulated receptor in the distal leg epithelium of digging *P. carolinus* (**Fig. 4b**). At the cellular level, we observed robust and specific epithelial localization of *t1r3* mRNA within the tips of papillae, which specify sensory leg function of digging sea robins (**Fig. 4c**). T1r3 is canonically expressed in the specialized receptor cells of oral tastebuds, where it forms obligate heterodimers with T1r1 or T1r2 to mediate sensation^7,13–15^ (**Extended Data 3a**). Indeed, we also detected expression of *t1r2* in papillae epithelial cells, albeit at lower levels, suggesting expression of functional heteromeric taste receptors (**Fig. 4c**). In contrast, we did not detect taste receptor expression in the legs of non-digging *P. evolans*, consistent with insensitivity to most common tastants (**Fig. 4d**). To determine whether the selectivity of sea robin taste receptors matches leg physiology and behavior, we heterologously expressed *P. carolinus* receptors in HEK293T cells and measured responses to chemical agonists. Remarkably, we found that co-expression of T1r2 and T1r3 mediated robust responses to the same L-amino acids that distinguish digging *P. carolinus* chemosensitivity from that of non-digging *P. evolans* (**Fig. 4e, Extended Data 3b,c**)^16,17^. Thus, papillae are molecularly defined polymodal sensory structures that specialize sea robin legs to support adaptive digging behavior.

**Figure 4.**
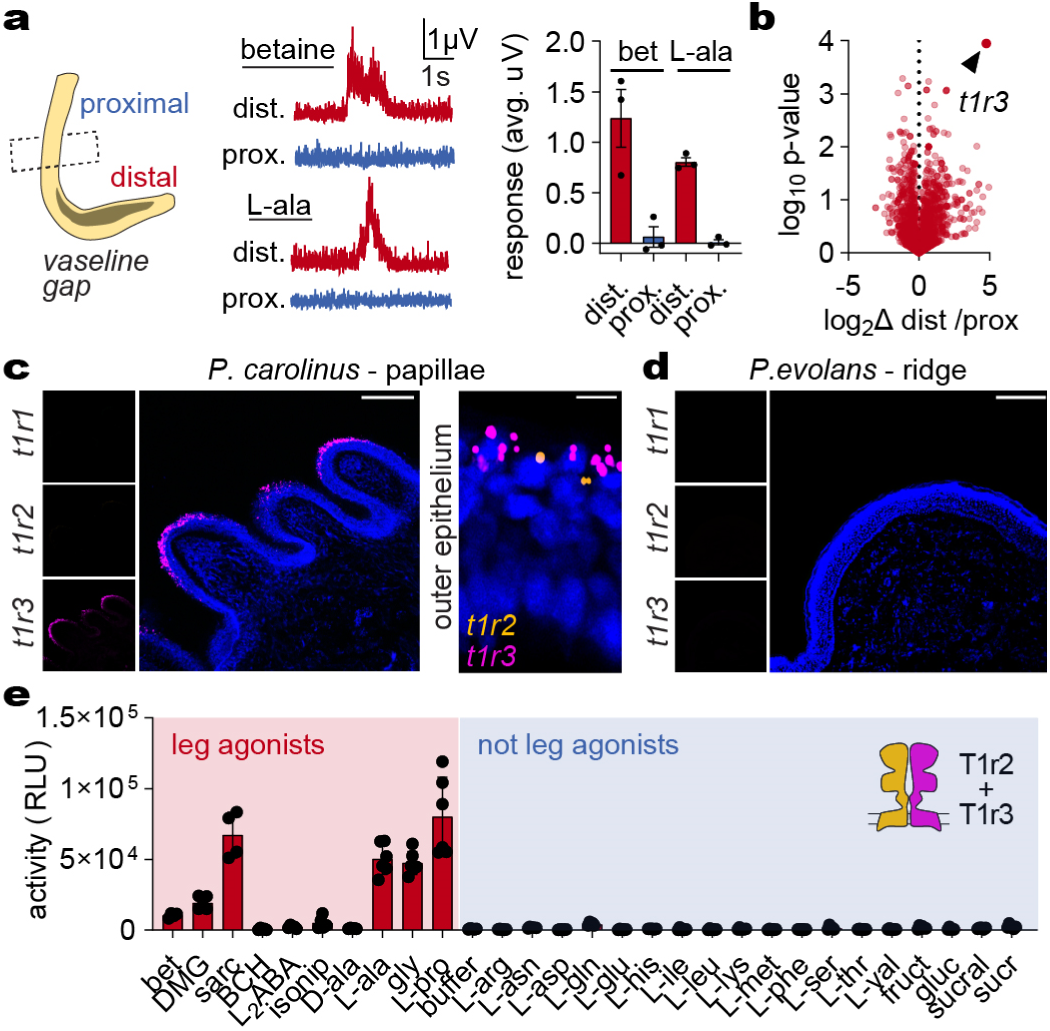
Taste receptors in papillae mediate leg chemosensation. **a**, Only papillae-covered distal legs, but not proximal legs, of digging *P. carolinus* responded to appetitive chemicals (2 mM betaine or L-alanine, n = 3, *p* < 0.0180 betaine, *p* < 0.0001 L-ala, t-test with Welch’s correction). **b**, The taste receptor *t1r3* (arrowhead) was the most enriched receptor in the chemosensitive distal leg epithelium. **c**, *t1r3* and *t1r2* were expressed in surface epithelial cells of papillae in digging *P. carolinus*, but absent from non-digging *P. evolans*, visualized by *in situ* hybridization (scale bars = 100 µm for *left, right*, 10 µm for *middle*, images representative of 3 animals per species). **d**, T1r2/T1r3 heterodimers (from *P. carolinus)* responded to the L-amino acids sensed by digging *P. carolinus* but not non-digging *P. evolans* (*n* = 6). RLU = Relative Light Units. Data in **b** and **e** represented as the mean ± s.e.m.

## Leg sensory organs as a major trait gain that enables novel behavior

To further interrogate the role of papillae in digging behavior we examined their function in three contexts: (1) sea robin ontogeny; (2) interspecies F1 hybrid animals; and 3) across the diverse phylogenetic tree of sea robins. Larval sea robins hatch without legs and are initially pelagic. Then, legs separate from pectoral fins during the first three weeks of development as the animals settle to the sea floor^18–20^ (**Fig. 5a**). While observing this transition we discovered that *P. carolinus* lacked papillae during the first five weeks of development (**Fig. 5b**). Strikingly, *P. carolinus* began to localize and dig up prey at a stage coinciding with papillae formation, suggesting that the presence of papillae facilitates digging behavior (**Fig. 5c**). Next, we considered the pattern of inheritance in crosses of hybrid animals with distinct parental traits. Crosses of digging *P. carolinus* males with non-digging *P. evolans* females produced viable offspring that could be raised through the stage of leg-separation (**Fig. 5d**). Hybrid animals developed legs that possess papillae which closely resemble the digging parent (**Fig. 5e**). Consistent with a critical role in sensation, papillae correlated with enhanced chemosensitivity and sensory digging behavior in hybrid sea robins (**Fig 5e-f**). These results collectively support the hypothesis that papillae are required for sensation and predatory digging.

**Figure 5.**
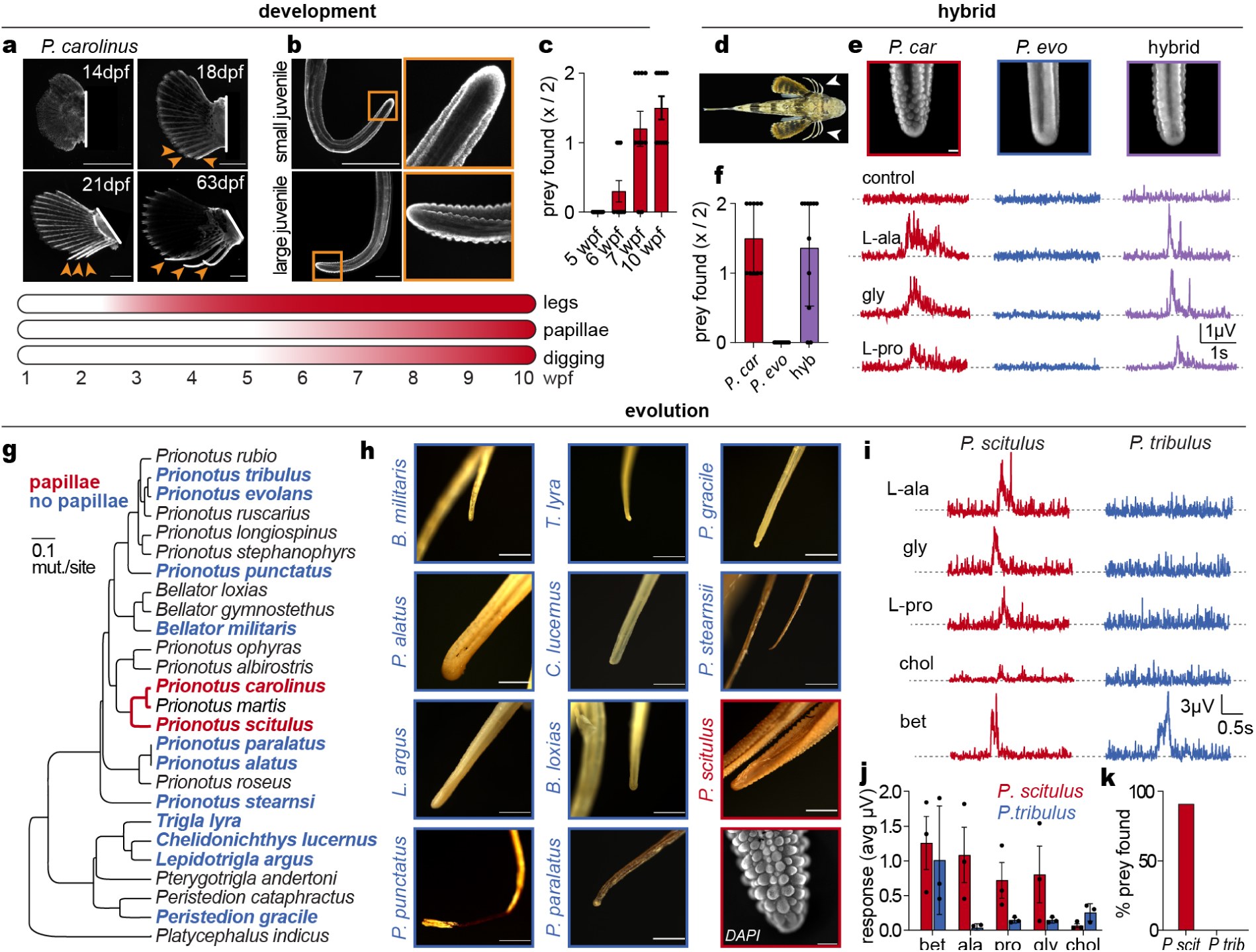
Papillae are a major gained trait and facilitate predatory digging behavior. **a**, *P. carolinus* legs formed weeks after hatching (stained with DAPI, scale bars = 1 mm). DPF = Days post fertilization. **b**, Legs of larval sea robins initially lacked papillae, which later formed in larger juveniles (scale bars = 1 mm). WPF = Weeks post fertilization. **c**, Onset of digging behavior correlated with papillae formation in developing *P. carolinus* sea robins (*n* = 10 trials per age group, key below indicates approximate timeline, ANOVA with Tukey’s post-hoc test: 5 vs 6 wk. n.s., 5 vs 7 wk. p < 0.0001, 5 vs 10 wk. p < 0.0001). **d**, Dorsal view of an F1 hybrid sea robin produced in a cross of a female *P. evolans* and male *P. carolinus.* **e**, Hybrid progeny have leg papillae (*top*, scale bar = 250µm) and responded to appetitive chemicals similar to their digging parent. Representative image of *n* = 4. **f**, Hybrid sea robins exhibited robust digging behavior to find prey, similar to parental *P. carolinus* that also have leg papillae (*n* = 10, *p* < 0.0001 hybrid or *P. carolinus* vs *P. evolans* by ANOVA with Tukey’s post-hoc test). **g**, Species with leg papillae (red) represent a restricted clade of the sea robin phylogenetic tree^30^. Species in blue lacked papillae, and species in black were not analyzed. **h**, Leg morphology of museum specimens and DAPI stained papillae in *P. scitulus* (scale bar = 200 µm). **i-j**, *P. scitulus*, which has leg papillae, responded to appetitive L-amino acids, while *P. tribulus*, which lacks papillae, was insensitive (*n* = 4 recordings per chemical per species). **k**, Papillae-expressing and chemosensitive *P. scitulus* exhibited robust digging behavior to find prey, while *P. tribulus* did not dig or find prey (*n* = 10 trials). Data in **c**, **f**, **j** represented as the mean ± s.e.m.

Finally, we leveraged the biodiversity of sea robin species from around the world to ask whether sensory specialization reflects the ancestral condition or is a recent innovation. We first used museum specimens to examine the leg morphology from a wide range of sea robin species (**Extended Data 4**). Surprisingly, we observed leg papillae in only a small group of sea robins comprising the closest relatives of digging *P. carolinus* (**Fig. 5g**, *red text*). In contrast, the legs of many species exhibited a stick-like morphology similar to fin rays, suggesting these structures may function as simple locomotory appendages that are ancestral to more advanced sensory legs (**Fig. 5h**). Guided by this analysis, we obtained and examined the physiology and behavior of two additional species, one with papillae (*Prionotus scitulus*) and one lacking papillae (*Prionotus tribulus*). Only *P. scitulus* exhibited chemosensory specialization and digging behavior, consistent with its leg morphology and close evolutionary proximity to *P. carolinus* (**Fig. 5i-k**). Therefore, leg papillae are sensory specializations unique to a specific clade of sea robin species and represent a novel trait gain.

We conclude that sensory digging behavior is restricted to a small lineage of specialist fishes that have diversified legs as novel sense organs, evolving unique molecular, morphological, and functional properties in sensory end-organs at the tips. We hypothesize that sea robins initially developed fin ray-like legs for locomotion. Ancestral organs then evolved limited sensory capability to facilitate manipulation of the visible substrate in search of food. Finally, evolution of sensory papillae further specialized legs to localize and uncover buried prey. Thus, sensory papillae represent a striking example of a major trait gain within a specific lineage that allows organismal expansion into a new behavioral niche. Indeed, our companion study demonstrates that legs use developmental programs that share features with other animal forelimbs and additionally contribute to the specialization of sensory structures (*Herbert, et al*). Future work will leverage the remarkable biodiversity of sea robins to understand the genetic basis of novel trait formation and diversification in vertebrates by focusing on molecular mechanisms of fin ray separation, adaptation of the nervous system, and acquisition of sensory properties. Thus, our work represents a basis for understanding how novel traits evolve to facilitate the lifestyles of diverse organisms across distinct ecologies.

## Acknowledgments

We thank: the Marine Biological Laboratory and Gulf Marine Specimens for providing and holding adult sea robins; A Grearson for photography; C Heacock and P Kilian for animal care; A Lee, W Valencia-Montoya, and the Harvard University Bauer Core for bioinformatics advice; E Soucy for assistance with electrophysiology; A Williston and the Harvard Museum of Comparative Zoology for loaning and assistance with museum samples; R Losick for comments on the manuscript; and the Harvard Medical School Electron Microscopy Core Facility. Roger Williams University for larval rearing and juvenile care; A Leonardi, J Sears, L Fitzgerald, A Grove Z. Sotero.

This research was further supported by grants to: NWB from the New York Stem Cell Foundation, Searle Scholars Program, and the NIH (R35GM142697); CAA from the Harvard Brain Initiative Postdoc Pioneer’s Award, The Charles A. King Trust Postdoctoral Fellowship, and the National Science Foundation (NSF-PRFB 2010728); ALH from a Helen Hay Whitney Fellowship, a Grass Fellowship in Neuroscience from the Marine Biological Laboratory, and three Early Career Whitman Fellowships from the Marine Biological Laboratory; D.M.K is an investigator of the Howard Hughes Medical Institute; MWB is supported by the Max Planck Society; AS from the European Research Council (ERC) under the European Union’s Horizon 2020 research and innovation programme (grant agreement No 101002724 RIDING), the Air Force Office of Scientific Research under award number FA8655-20-1-7028, and the National Institutes of Health (NIH) under award number R01DC018789.

## Author Contributions

CAHA, SPK, BLW, NWB contributed to molecular, cellular, and organismal studies. ALH and DMK contributed to developmental and hybrid studies. AS performed all mathematical modelling. QL and MWB performed T1r functional assays. ALR and ANG established animal culture methods. All authors were involved with writing or reviewing the manuscript.

## Methods

### Animals

Adult wild-caught sea robins (*Prionotus carolinus* and *Prionotus evolans*) were provided by the Marine Biological Laboratory, Woods Hole, MA, and kept on a 12 hr light/dark cycle in natural sea water. Larval sea robins were provided by Dr. Amy Herbert (see companion article), fed daily, and kept on a 12 hr light/dark cycle in natural sea water. Where required for experiments, animals were euthanized by immersion in Tricaine-S (Syndel) until 10 minutes past cessation of opercular movements. Animal protocols were approved by the Harvard University and Roger Williams University Animal Care and Use Committees (protocols HU: ID 18-05-324-1, RWU: R19-07-09).

### Behavior experiments

#### Adult behavior

Sea robins were placed in a 1.5 m diameter pool containing a volume of ∼750 L (or 200 gallons) of sea water and ∼8 cm of sand. Experiments were filmed from above by a GoPro camera and evaluated using Windows Media Player (Microsoft). Fish were acclimated for 20 minutes in the behavior tank before experiments. Prey items were added to the tank at indicated depths just prior to experiment start using an opaque divider to prevent visualization of the bury site. Mussels were opened and halved along the hinge. Capsules were placed inside of a cleaned mussel shell to provide a physical object for capture. All behavior experiments were run for 30 minutes, and scored as negative if the mussel was not found within this time. Trials were scored as successful if prey items were excavated, or if a characteristic “head snap” behavior was noted at the site of buried prey. For experiments involving much smaller *P. scitulus* and *P. tribulus* species, smaller tanks and mussels were used (9 L, 23 x 34 cm).

#### Juvenile behavior

Groups of juvenile sea robins (*P. carolinus*, *P. evolans*, and hybrids) were acquired from Roger Williams University and raised in seawater tanks. Digging behavior experiments were performed using a behavioral model first tested with adult sea robins. Prior to behavioral experiments, sand was thoroughly cleaned, and animals were not fed that morning.

Blue mussels (*Mytilus edulis*) were purchased from the local market and dissected into 1-1.5 cm pieces for shallow sand burial. Two PVC-pipe pieces were used to shield visual cues while two mussel pieces were separately buried in the sand. The PVC-pipes were removed, allowing animals to explore the sand. Trials were filmed with GoPro HERO 7 cameras (GoPro, Inc.), propped outside of the behavioral tank. Trials were run for 30 minutes, then the tanks were reset for a new trial by removing any excess mussel pieces. The mussel pieces changed position in the tank each time.

GoPro videos were uploaded and played back in Windows Media Player to assess capture success. Capture success was defined as a fish locating the mussel, evidenced by mussel consumption or a ‘head snap’ where a fish opens its mouth and snaps in the direction of the prey item. Each group tank had two mussel pieces or ‘chances of success’ per trial. Data was compiled and visualized in Prism GraphPad.

### Molecular Biology

Complete cDNA sequences of taste receptors were obtained using a SMARTer 5’/3’ kit for RACE PCR to obtain 5’ sequence information missing from reference transcriptome (Takara, #634858). Briefly, RNA was extracted from oral and leg tissues using standard Trizol extraction, purified with Zymo Clean and Concentrator Kit (Zymo research #R1013), and diluted in 10-20 mL RNase free H2O without DNAase treatment. Taste receptor specific primers were used to amplify the 5’ end and PCR products were cloned as suggested by the company and verified by sequencing:

*t1r1* 5’ RACE primer: 5’-GATTACGCCAAGCTTGGCAGTGGTATTCATTTGTGGCCTCAGG
*t1r2* 3’ RACE primer: 5’-GATTACGCCAAGCTTCATGATCTTGCTTCTCTGTCTCTGCTGG
*t1r3* 5’ RACE primer: 5’-GATTACGCCAAGCTTTTAGTTCCCCGGCTCTGGCTGTGGCTGT

All constructs were codon-optimized and synthesized by Genscript (Piscataway, NJ) into the pEAK10 expression vector.

### Mathematical modeling

Chemical cues from a buried target spread mostly by diffusion because in sand water flow is largely suppressed. Qualitatively, chemicals diffuse over a spatial scale *l* that depends on the time *t* since it was released from the target 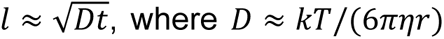 is the chemical diffusivity and depends on the temperature and viscosity of water (*T* and *η*) and the size of the odorant molecule (*r*). Leg agonists are small molecules and their diffusivity in water at room temperature is typically *D* ≈ 10^−10^ to 10^−9^*m*^2^/*s* (*D* ≈ 5 × 10^−10^*m*^2^/*s* for dopamine^21^ and fluorescein^22^ and closer to *D* ≈ 10^−9^*m*^2^/*s* for various aminoacids^23^). Thus, by the end of our experiments, the chemicals may only travel 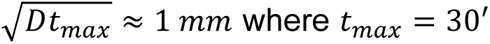 is the duration of the experiment. This simple argument shows that only a thin ≈ *mm* layer of sand in contact with the mussel is imbued with chemical signal, hence fish must touch sand close to the mussel in order to detect a signal.

In order to move beyond this qualitative argument, we focus on the betaine-capsule experiment for which we can predict the odor concentration everywhere in space and time and compare it with sensitivity by the legs. Betaine is a small amino acid derivative and we control its volume and concentration in the capsule (*c*_0_ = 0.1 mM; 1 mM; 10 mM or 100 mM; *V* = 1 *m*L; thus the capsule contains *M* = *c*_0_*V* moles of betaine; we use *D* ≈ 10^−9^*m*^2^/*s* relevant for aminoacids^23^). Additionally, we know that the legs sense betaine above a threshold concentration *c*^∗^ = 10 to 100 μ*M* **(Fig. 2e)**.

To complete the mathematical model, we need boundary conditions. The boundaries of the tank are irrelevant since they are much further than the diffusive length scale ≈ 1 *mm*. The boundary condition at the water/sand interface depends on flow of water in the tank above sand. Since we do not control water flow in the experiments, we consider two opposite scenarios; the actual dynamics is somewhere in between these scenarios. If water is perfectly still, there is no boundary condition at the sand/water interface. A small portion of the capsule located at *r*_*ss*_ = (*x*_*s*_, *y*_*s*_, *z*_*s*_) releases *dM* = *Mdx*_*s*_*dy*_*s*_*dz*_*s*_ moles of betaine and results in a molar concentration of betaine *dc* at any point in space and time^24^:

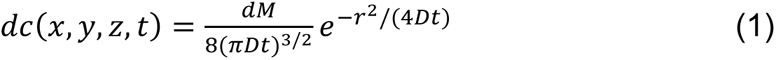

where *r*^2^ = (*x* − *x*_*s*_)^2^ + (*y* − *y*_*s*_)^2^ + (*z* − *z*_*s*_)^2^; the water/sand interface is located at *z* = 0, water is at *z* < 0 and sand is at *z* > 0. Second, if water flows fast, betaine is greatly diluted away resulting in a Dirichlet boundary condition and using the method of images we obtain a modified prediction:

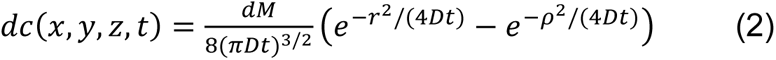

where ρ^2^ = (*x* − *x*_*s*_)^2^ + (*y* − *y*_*s*_)^2^ + (−*z* − *z*_*s*_)^2^. The total betaine concentration is obtained in both scenarios by summing the contribution *ddcc* from all portions of the capsule. Integrating over the volume *VV* of the capsule (a prolate ellipsoid of dimensions 6.3 mm and 16 mm located at *zz* = *l*) we obtain:

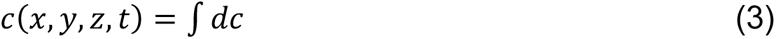

We compute the integral in equation (3) numerically. The concentration of betaine in the topmost ≈ 0.5 *mmmm* layer of sand, presumably sampled by legs **(Fig 1h)**: the two scenarios provide similar predictions hence we will ignore water flow for our current purposes. The conclusion from this calculation is that the concentration of betaine in sand is initially too small to be detected by the legs. After an initial transient, betaine can be detected on sand above the capsule. The duration of the initial transient depends on betaine concentration in the capsule, consistent with experiments in **(Fig. 1i)** showing fish finds most efficiently the more concentrated capsules.

### Electron microscopy

Electron microscopy was performed by the Harvard Medical School Electron Microscopy Core Facility according to standard protocols. Briefly:

#### Scanning electron microscopy

Samples were fixed overnight in a mixture of 1.25% formaldehyde, 2.5% glutaraldehyde and 0.03% picric acid in 0.1 M Sodium cacodylate buffer, pH 7.4, washed with 0.1 M sodium cacodylate buffer, and post fixed with 1% osmium tetroxide in 0.1 M sodium cacodylate buffer for 2 hours. Tissues were then rinsed in ddH_2_0 and dehydrated through an ethanol series (30%, 50%, 70%, 95%, (2x)100%) for 15 min per solution. Dehydrated tissues were dried in an autosamdri-815 critical point dryer and mounted on aluminum stages with silver paint and coated with platinum (10 nm). The dried tissues were imaged on a Hitachi S-4700 Field Emission Scanning Electron Microscope (FE-SEM) at an accelerating voltage of 5kV.

#### Transmission electron microscopy

Samples were fixed overnight in fixative solution (1.25% formaldehyde, 2.5 % glutaraldehyde, and 0.03% picric acid in 0.1 M sodium cacodylate buffer, pH 7.4), then washed with 0.1 M sodium cacodylate buffer and post-fixed with 1% osmium tetroxide/1.5% potassium ferrocyanide (in H_2_O) for 2 hours. Samples were then washed in a maleate buffer and post fixed in 1% uranyl acetate in maleate buffer for 1 hour. Tissues were rinsed in ddH_2_0 and dehydrated through an ethanol series (50%, 70%, 95%, (2x)100%) for 15 min per solution. Dehydrated tissues were put in propylene oxide for 5 min before they were infiltrated in epon mixed 1:1 with propylene oxide overnight at 4 °C. Samples were polymerized in a 60 °C oven in epon resin for 48 hours. They were then sectioned into 80 nm thin sections and imaged on a JEOL 1200EX Transmission Electron Microscope.

### Transcriptomics

We generated tissue-specific *de novo* transcriptomes for *Prionotus carolinus* and *Prionotus evolans* (Supplementary Table 1). Tissues were dissected and stored frozen in RNAlater until use. RNA extraction, library preparation, and RNA sequencing were performed by the Harvard Bauer Core Facility using NovaSeq (2×250) platform or by Genewiz (Azenta) using a HiSeq (2×150 bp) platform (see Supplementary Table 1). Adaptor trimming was performed using trimgalore^25^, and reference transcriptomes were assembled *de novo* using Trinity^26^. Open reading frames were identified using transdecoder^26^. Reads were pseudo-aligned and transcript abundance was estimated using Kallisto^27^ and our novel transcriptome assemblies as a reference. Annotation was performed using Diamond^28^.

### Fluorescence In situ hybridization (RNAscope)

Tissues collected from recently euthanized specimens were immediately frozen in OCT and stored at - 80°C until use. Blocks were trimmed and sectioned (18-50 µm) using a cryostat (Leica CM3050S). Probes were designed by ACD Inc. and the manufacturer-recommend protocol was followed as described for fresh frozen tissues. Pretreat 3 was used for 30min and fluorescent probes used included TSA-FITC, TSACy3 and TSA-Cy5 (Perkin Elmer #NEL744E001KT and #NEL754001KT), samples were mounted in ProLong Gold (Thermo Fisher #P36931) with DAPI, and imaged with a LSM 980 Confocal Microscope with Airyscan2 (Zeiss), LSM 900 Confocal Microscope (Zeiss), or AxioZoom v16 microscope. Images were processed using FIJI (NIH).

### Primary neuron culture

Primary neuron culture was based on protocols established for zebrafish^29^ and optimized to improve cell survival as determined by cell density, neurite out-growth, and expression of voltage-gated conductances. Animals were euthanized and spinal ganglia corresponding to the legs and/or pectoral fin were dissected and placed in ice-cold HBS (HEPES buffered saline). Special care was taken to remove blood from the tissue, as blood cells persist through the cell purification protocol. Ganglia were cut into small pieces and cells were dissociated by incubating ganglia pieces in enzyme solution (10 mg/mL collagenase IV (Worthington), in 1XHBS (HEPES buffered saline, Thermo) for 1 hour at room temperature with gentle agitation. Enzyme solution was carefully removed and replaced with 10 mL culture media (L-15 media (Gibco #11415064) supplemented with 10% fetal calf serum (FBS) (Gibco) and 50 I.U./mL penicillin and 50 μg/mL streptomycin (Gibco), and 0.5 nM neurotrophin-3 (NT-3, Sigma #SRP6007). Cells were released by trituration through fire-polished glass pipettes of decreasing diameter, until tissue was nearly completely broken up. Undissociated tissue was removed by passing cells through 100 µm nylon mesh filters (Corning). Debris and glial cells were removed by centrifuging cells over a 2 mL 15% bovine serum albumin (BSA, Sigma) solution in culture media at 250xg for 10 minutes at 4°C. The entire supernatant was removed and the pellet containing enriched sensory neurons was resuspended in fresh 1 mL culture media. For plating, glass-bottomed Mattek dishes coated with poly-L-lysine were further coated in 1 mg/mL laminin in H_2_0 (Corning) for 1 hour, and rinsed 3X with sterile water. 100 µL cells were spotted into the center of each dish, allowed to adhere for several hours before addition of 1 mL of culture media. Cells were cultured in humidified chambers at 17°C for 3-5 days before use in functional experiments.

### Calcium imaging

Cultured neurons were washed in extracellular solution (in mM): 140 NaCl, 5 KCl, 10 HEPES, 2 CaCl2, 2 MgCl2, 10 D-galactose pH 7.4. and loaded with 5µM Fluo-8 AM (Abcam) and 0.01% Pluronic F-127 diluted in extracellular solution for 15min in the dark at room temperature, and washed 3X in extracellular solution before imaging. Imaging experiments were performed using a custom inverted wide-field fluorescence microscope running Metamorph imaging software. Perfusion experiments were performed using a SmartSquirt Micro-Perfusion system (Automate Scientific) pressurized to ∼30 kPa. Stimuli were delivered for one minute followed by two minute wash intervals.

### Museum Samples

Museums samples (Supplemental Table 2) were staged in ethanol in glass dishes and legs were imaged using an AxioZoomV16 microscope equipped with a color camera (Zeiss). Images of entire animals were taken with a Cybershot digital camera (Sony).

### Cell culture

HEK293 cells (ATCC, authenticated and validated as negative for mycoplasma by vendor) were cultured in Dulbecco’s Modified Eagle Medium (DMEM) (Gibco) supplemented with 10% fetal calf serum (FBS) (Gibco) and 50 I.U./mL penicillin and 50 μg/mL streptomycin (Gibco) at 37 °C, 5% CO_2_ using standard techniques. For transfection, HEK293 cells were washed with Opti-MEM Reduced Serum Media (Gibco) and incubated with transfection mix containing 1 µg total of the indicated plasmid DNA and 3 µL Lipofectamine 3000 Transfection Reagent (Invitrogen) in Opti-MEM for 4-8 hours at 37 °C. Cells were then replated onto glass coverslips in DMEM, incubated for 2 hr at 37 °C, and then moved to 30 °C incubation overnight.

### Histology

#### Whole Mount Histology

Tissues were fixed in 4% paraformaldehyde (PFA) in PBS for 5 hr on a rocker and subsequently stored in PBS at 4°C until use. For immunofluorescence, samples were then washed with PBST (Triton X-100, 0.1%) 3 times and blocked for 1 hr in 10% normal goat serum (NGS) in PBST and antibody solution containing anti-HNK1 polyclonal antibody (1:100) (1C10 deposited by Halfter, WM Developmental Studies Hybridoma Bank) and anti-calretinin (1:100) (clone 3k22 #ZRB5054, Sigma) was applied for 24 hr at 4°C in NGS. Samples were then washed with PBST (TritonX 0.1%) 3 times, and goat anti-rabbit IgG H&L (Alexa Fluor® 488) (Abcam ab6702) and Goat Anti-mouse IgG H&L (Cy3 ®) (AbCam ab97035) secondary antibodies were added (1:500). Finally, the samples were washed 3-5 times in PBST, mounted in Vectashield (Vector Laboratories), and imaged with an AxioZoom V16 Zoom Microscope (Zeiss) or LSM 980 Confocal Microscope with Airyscan2 (Zeiss). For DAPI staining, tissues were stained in 1µg/mL DAPI in 1XPBS for 30 min, washed 3X in PBS, and staged for imaging. Images were processed using FIJI.

#### Immunofluorescence staining of HEK Cells

HEK293T cells were transfected as described above. Cells were washed 3 times in PBS and fixed in 4% paraformaldehyde (PFA) in PBS for 10 min at room temperature. Coverslips were then incubated in blocking mixture containing 1% bovine serum albumin (BSA) and 10 mM glycine in PBS. Samples were then incubated in antibody solution containing anti-FLAG®M2 monoclonal antibody (Sigma F1804) (1:100) and HA tag Polyclonal Antibody (Proteintech ab51064-2-AP) (1:250) for 24 hr at 4 °C in 1% BSA. Samples were then washed with PBS 3 times, and secondaries antibody mix containing goat anti-rabbit IgG H&L (Alexa Fluor® 488) (Abcam ab6702) (1:500) and Goat Anti-mouse IgG H&L (Cy3 ®) (AbCam ab97035) (1:500) was added. Finally, the samples were washed 3 times in PBS, mounted in Vectashield with DAPI (Vector Laboratories), and imaged with a LSM 980 Confocal Microscope with Airyscan2 (Zeiss). Images were processed using FIJI.

#### Thin Section

Tissues from the indicated species were fixed in 4% paraformaldehyde (PFA) in PBS for approximately 5 hr on a rocker and dissected into small sections, washed with PBS 3 times, and incubated in 30% sucrose in PBS at 4 °C on ice. After embedding in OCT (optimal cutting temperature compound), samples were frozen and sectioned using a cryostat (Leica CM3050S) at 20-50 µm sections. After subsequent washes in PBST, samples were blocked for 1 hr in 5% normal goat serum (NGS) in PBST and antibody solution containing anti-HNK1 polyclonal antibody (1:100) (1C10 deposited by Halfter, WM Developmental Studies Hybridoma Bank) and anti-calretinin (1:100) (clone 3k22 #ZRB5054, Sigma) in PBS was applied for 24 hr at 4 °C. Samples were then washed with PBST (TritonX 0.1%) 3 times, and goat anti-rabbit IgG H&L (Alexa Fluor® 488) (Abcam ab6702) and Goat Anti-mouse IgG H&L (Cy3 ®) (AbCam ab97035) (1:500) secondary antibodies were added (1:500). Finally, the samples were washed 3-5 times in PBST, mounted in Vectashield with DAPI (Vector Laboratories), and imaged with a LSM 980 Confocal Microscope with Airyscan2 (Zeiss). Images were processed using FIJI.

### T1r functional assays

Sea robin T1r plasmids were purified (Qiagen Plasmid Mini Kit) and *in vitro* responses were measured using a luminescence cell-based assay^16,17^. HEK293T cells (from the Matsunami Laboratory, Duke University, USA) were transiently co-transfected with expression vectors of sea robin T1rs, a rat G-protein (Gα15i2) and a photoprotein (mt-apoclytin-II) using Lipofectamine 2000 (Invitrogen). Control cells were only transfected with plasmids for the rat G-protein and the photoprotein (without receptor plasmids). Two days after transfection, cells were seeded in 96-well black-walled CellBIND plates (Corning) and assayed on a FlexStation3 microplate reader (Molecular Devices), which applied ligands and measured the luminescence intensity over 110 seconds in each well. All ligand solutions of various concentrations were prepared in filtered HEPES solution (4-(2-hydroxyethyl)-1-piperazine ethanesulfonic acid) at pH 7.4 (sugars: 100 mM, n = 6; L-amino acids: 50 mM, n = 6; D-alanine, isonipecotic acid, N,N-dimethylglycine, and L-2-aminobutyric acid: 50 mM, n = 5; sarcosine: 50 mM, n = 3; betaine and 2-amino-2-norbornanecarboxylic acid: 12.5 mM, n = 5; no ligand buffer, n = 6). The ligand-evoked responses were quantified as the area under the curve (AUC) and expressed as relative light units (RLU).

### Patch clamp electrophysiology

Patch clamp recordings were carried out at room temperature using a MultiClamp 700B amplifier (Axon Instruments) and digitized using a Digidata 1550B (Axon Instruments) interface and pClamp software (Axon Instruments). Whole-cell recording data were filtered at 1 kHz and sampled at 10 kHz. Pipette resistances were 3-5 MΩ. The standard extracellular solution contained (in mM): 140 NaCl, 5 KCl, 10 HEPES, 2 CaCl_2_, 2 MgCl_2_, pH 7.4. The intracellular solution contained (mM): 140 Cs^+^ methanesulfonate, 1 MgCl_2_, 5 NaCl, 10 CsEGTA, 10 HEPES, 10 sucrose, pH 7.2. Whole-cell recordings were used to assess mechanical sensation with a piezoelectric-driven (Physik Instrumente) fire-polished glass pipette (tip diameter ∼1 μm). Mechanical steps 150ms in duration were applied in 1 μm increments were applied every 5 s while cells were voltage-clamped at −80 mV.

### Nerve recording

Legs from sedated animals were transferred to a 10 cm petri dish containing 50 mL holding solution (430 mM NaCl, 10 mM KCl, 10 mM HEPES, 10 mM CaCl_2_, 50 mM MgCl_2_, 10 mM D-glucose, pH 7.6). Nerve recordings were performed using a borosilicate glass suction electrode shaped and polished to fit over the entire cut end of the leg nerve. A similar reference electrode was placed in the bath. Gap-free recordings were made with 10 kHz sampling at 10,000 x gain, and signals were highpass filtered at 100 Hz and lowpass filtered at 1 kHz using a Warner DP-311A headstage and AC/DC amplifier (Warner), and digitized using a Digidata 1440A digitizer (Molecular Devices) using ClampEx software (Molecular Devices). Recordings were processed using Clampfit (Molecular Devices). For quantification of responses, the absolute value of the signal was processed using a lowpass 25 Hz gaussian filter, the baseline signal was subtracted, and the response amplitude was integrated over the response area. Transients from pipette-bath-contact were removed from figure traces and not quantified. For controls and stimuli where no response was observed, a similar response area was measured as for leg agonists. For experiments testing chemical sensitivity of proximal versus distal legs, chambers were drawn around each leg region using blunted syringes loaded with 5% mineral oil mixed with Vaseline to create water-tight boundaries, and chemicals were perfused into the indicated chamber.

### Agonists

Chemicals screened included [Chemical name, (Abbreviation, source)]: betaine (Bet, Sigma-Aldrich #61962, dimethylglycine (DMG, Sigma-Aldrich #D1156), sarcosine (sarc, Sigma-Aldrich #131776), isonipecotic acid (isonip, Sigma-Aldrich #I18008), β-alanine (β-ala, Sigma-Aldrich #146064), nipecotic acid (nip. Sigma-Aldrich #211672), L-pipecolic acid (pip, Sigma-Aldrich #P2519), у-aminobutyric acid (GABA, Hello Bio #HB0882), L-2-aminobutyric acid (L_2_ABA Sigma-Aldrich #A1879), trimethylamine hydrochloride (TMA, Sigma-Aldrich #T72761), serotonin hydrochloride (5-HT, Sigma-Aldrich #H9523), tyramine (tyrmn, Sigma-Aldrich T90344), L-carnitine hydrochloride (L-carn, Sigma-Aldrich #C0283), taurine (taur, Sigma-Aldrich T0625), L-proline (L-pro, Sigma-Aldrich P0380), Glycine (gly, Sigma-Aldrich #410225), L-cysteine (L-cys, Sigma-Aldrich #168149), L-alanine (L-ala, Sigma-Aldrich #W381829), L-leucine (L-leu,Sigma-Aldrich #L8000), L-tyrosine (L-tyr, Sigma-Aldrich #T3754), L-phenylalanine (L-phe, Sigma-Aldrich #P2126), L-glutamine (L-gln, Sigma-Aldrich #G3126), L-methionine (L-met, Sigma-Aldrich #M22060), L-aspartic acid (L-asp, Sigma-Aldrich #A9256), L-histidine (L-his, Sigma-Aldrich #H8125), L-ornithine (L-orn, Sigma-Aldrich #O2375), L-tryptophan (L-trp, Sigma-Aldrich #T0254), L-arginine (L-arg, Sigma-Aldrich #A5006), L-lysine (L-lys, Sigma-Aldrich #L5501), L-glutamate (L-glu, Sigma-Aldrich #G1251), L-serine (L-ser, Sigma-Aldrich #S4500), D-alanine (D-ala, Sigma-Aldrich #A7377), D-cysteine (D-cys, Sigma-Aldrich #30095), D-2-aminobutyric acid (D_2_ABA, Sigma-Aldrich #116122), D-serine (D-ser, Sigma-Aldrich #S4250), D-leucine (D-leu, Sigma-Aldrich #855448), D-cycloserine (DCS, Sigma-Aldrich #C6880), D-(+)-Trehalose dihydrate (treh, Sigma-Aldrich #T9531), Sucrose (sucr, Sigma-Aldrich #S7903), D-(+)-Galactose (D-gal, Sigma-Aldrich #G6404), D-(+)-Glucose (D-gluc, Sigma-Aldrich #49152), Eucalyptol (euc, Sigma-Aldrich #C80601), (R)-(+)-Limonene (lim, Sigma-Aldrich #183164), polygodial (polyg, Cayman #14979), carvacrol (carv, Sigma-Aldrich #W224502), nootkatone (nootk, Sigma-Aldrich #W316620), sodium acetate trihydrate (acet, Sigma-Aldrich #236500), sodium butyrate (butyr, Sigma-Aldrich #B5887), sodium propionate (prop, Sigma-Aldrich #P1880), chloroquine diphosphate (chlor, Sigma-Aldrich #C6628), denatonium benzoate (den, Sigma-Aldrich #D5765), Benztropine mesylate (benz, Sigma-Aldrich SML0847), Gaboxadol hydrochloride (gabox, Sigma-Aldrich #T101), α-(Methylamino)isobutyric acid (MeAIB, Sigma-Aldrich #M2383), BCH (BCH, Tocris #5027), Adenosine (aden, Sigma-Aldrich #A9251), Inosine (inos, Sigma-Aldrich #I4125), Acetylcholine chloride (ach, Sigma-Aldrich #A6625), Isoamyl acetate (isoam, Sigma-Aldrich #W205532), Choline chloride (chol, Sigma-Aldrich #C7527), Yoda1 (Selleck, S6678). Insoluble chemicals were first reconstituted in DMSO, and then diluted in ND96 to 0.1%-1% final DMSO. Fish extract was prepared by blending 1g frozen zebrafish per 2mL of holding solution, followed by centrifugation at 3,000 x g for 15 minutes to remove debris.

### Quantification and statistical analysis

Data were analyzed with Clampfit (Axon Instruments), Prism Graphpad and represented as mean ± SEM. N represented independent experiments for the number of cells/patches or behavioral trials. Data were considered significant if p < 0.05 using paired or unpaired two-tailed Student’s t tests, Wilcox test, or one- or two-way ANOVAs. All significance tests were justified considering the experimental design and we assumed normal distribution and variance, as is common for similar experiments. Sample sizes were chosen based on the number of independent experiments required for statistical significance and technical feasibility.

### Data Availability

Sequencing data will be uploaded to the Sequence Read Archive (SRA) repository and GenBank upon publication.

---

**Extended Data Figure 1.**
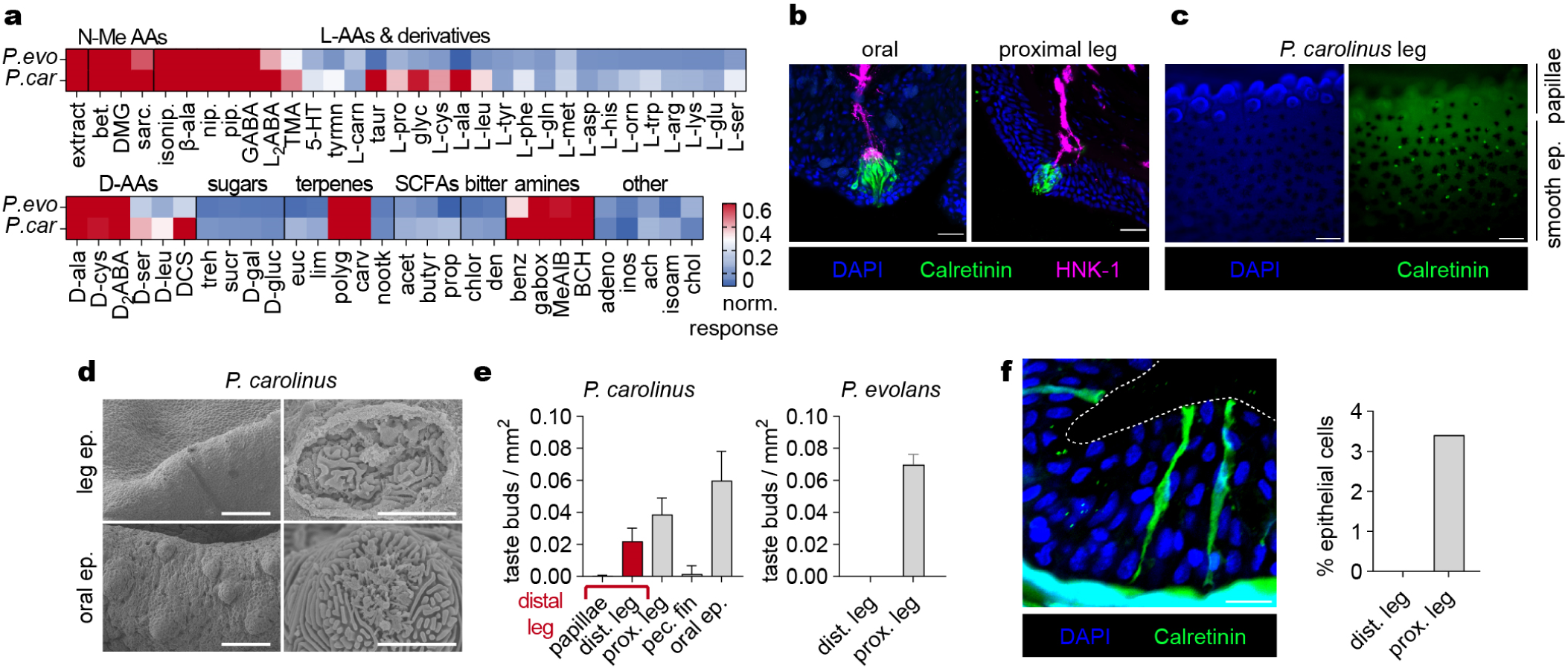
Sensory profiling of sea robin legs. **a**, Chemosensory responses from digging *P. carolinus* and non-digging *P. evolans* legs. Only *P. carolinus* were sensitive to appetitive L-amino acids. Heatmap of relative responses from > 6 legs. **b**, Proximal legs of *P. carolinus* express calretinin in spinal-shaped cells that resemble taste buds which are commonly found in fish skin^15^. Scale bars = 20 µm (left) and 200 µm (right). **c-e**, These structures were absent from papillae as analyzed by immunofluorescence and scanning electron microscopy. SEM scale bars 100 µm (left) and 5 µm (right). Data represented as the mean ± s.e.m. **f**, Calretinin-positive cells resembling solitary chemosensory cells were sparsely distributed throughout the skin of the proximal leg but absent from papillae of *P. carolinus*. Scale bar = 10 µm.

**Extended Data Figure 2.**
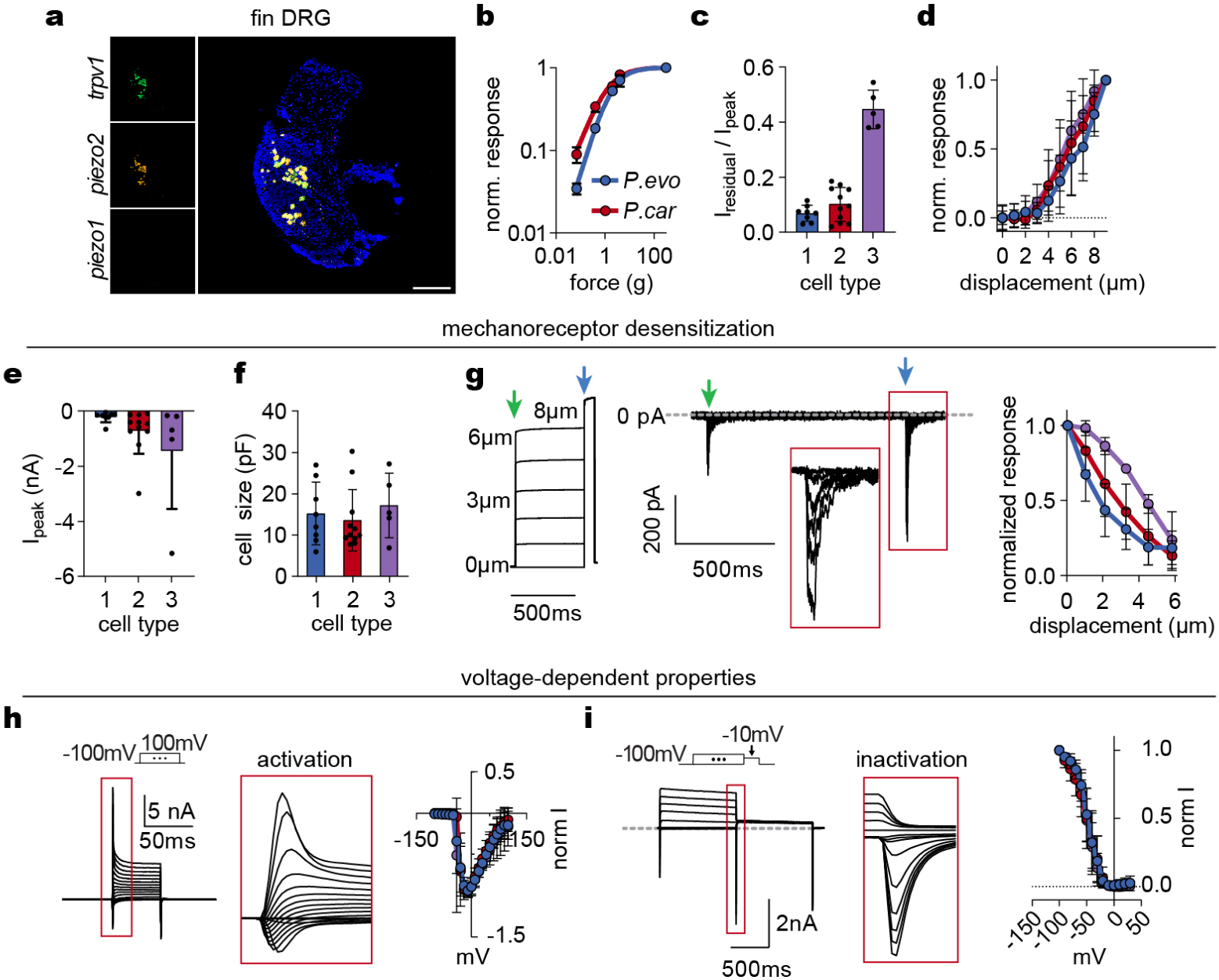
Mechanosensory properties. **a**, Fin-specific spinal ganglia contained *trpv1*-positive sensory neurons (green) that expressed the mechanoreceptor *piezo2* (orange) but not *piezo1* (pink, DAPI in blue, scale bar = 200 µm). Representative images of 3 animals per species. **b**, Force-response curve using filament stiffness shows higher sensitivity at low force for digging species *P. carolinus* compared with *P. evolans (*n > 5 stimulus-induced responses in recordings from 3 different legs). **c**, Desensitization kinetics as measured by “residual” current at the end of the mechanical stimulus over peak stimulus-induced currents. Responsive cells could be segregated into three distinct mechanosensitive neuron populations (*n* = 26, 8 type 1 with fast desensitization, 11 type 2 with intermediate desensitization, and 5 type 3 with slow desensitization). **d**, Mechanosensitive neuron populations exhibited similar thresholds for mechanosensitive responses (n = 8 type 1 in blue, 11 type 2 in red, 5 type 3 in purple). **e**, Peak mechanosensitive currents in three mechanosensitive neuron subtypes (*n* = 8 type 1, 11 type 2, 5 type 3). **f**, Capacitance of mechanosensitive neuron subtypes indicating approximate cell size (*n* = 8 type 1, 11 type 2, 5 type 3). **g**, Desensitization of mechanically-induced currents in response to increasing long duration displacement was similar across subtypes (*n* = 8 type 1, 11 type 2, 5 type 3). **h-i**, Voltage-dependent activation and inactivation of inward currents was similar across mechanosensitive neuron subtypes (n = 7 type 1, 10 type 2, 3 type 3). Data in **b**-**i** represented as the mean ± s.e.m.

**Extended Data Figure 3.**
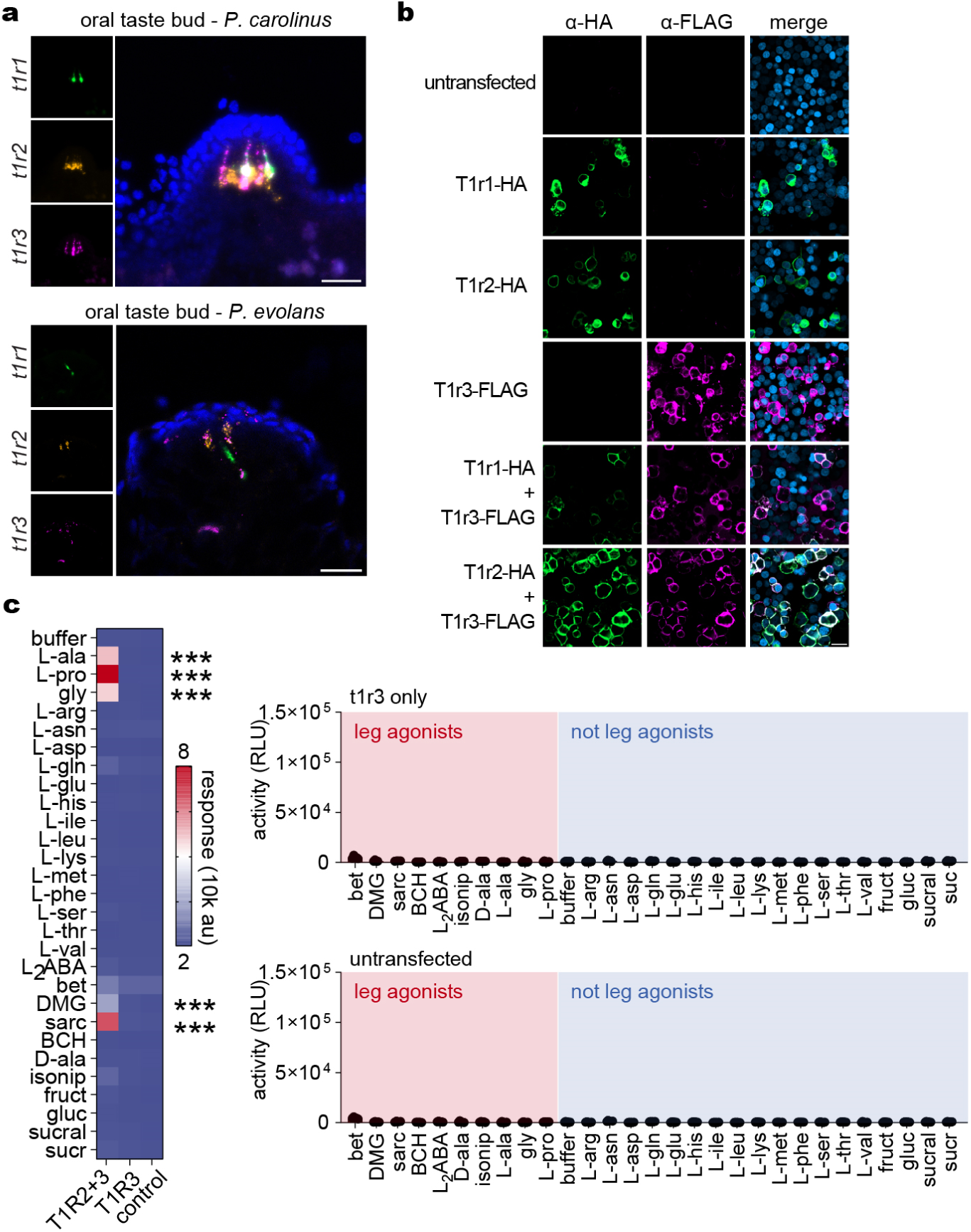
Taste receptor expression and function. **a**, *t1r1*, *2*, and *3* were expressed in oral taste buds of *P. carolinus* and *P. evolans*, visualized by *in situ* hybridization (scale bars = 25 µm, images representative of 3 animals per species). **b**, Epitope-tagged and codon optimized taste receptors were robustly expressed and trafficked to the plasma membrane in transfected HEK293T cells, as visualized by immunofluorescence microscopy. Scale bar = 25 µm. **c**, Control HEK293T cells (lacking T1R receptors) and cells expressing only T1R3 did not respond to L-amino acids or other molecules like T1R2-T1R3 heterodimers (n = 6, two-way ANOVA with Tukey’s post-hoc test. RLU = Relative Light Units. *** indicates *p* < 0.0001. Statistical significance indicates comparison of T1r2 + T1r3 co-expression to T1r3 only and control cells. Data represented as the mean ± s.e.m.

**Extended Data Figure 4.**
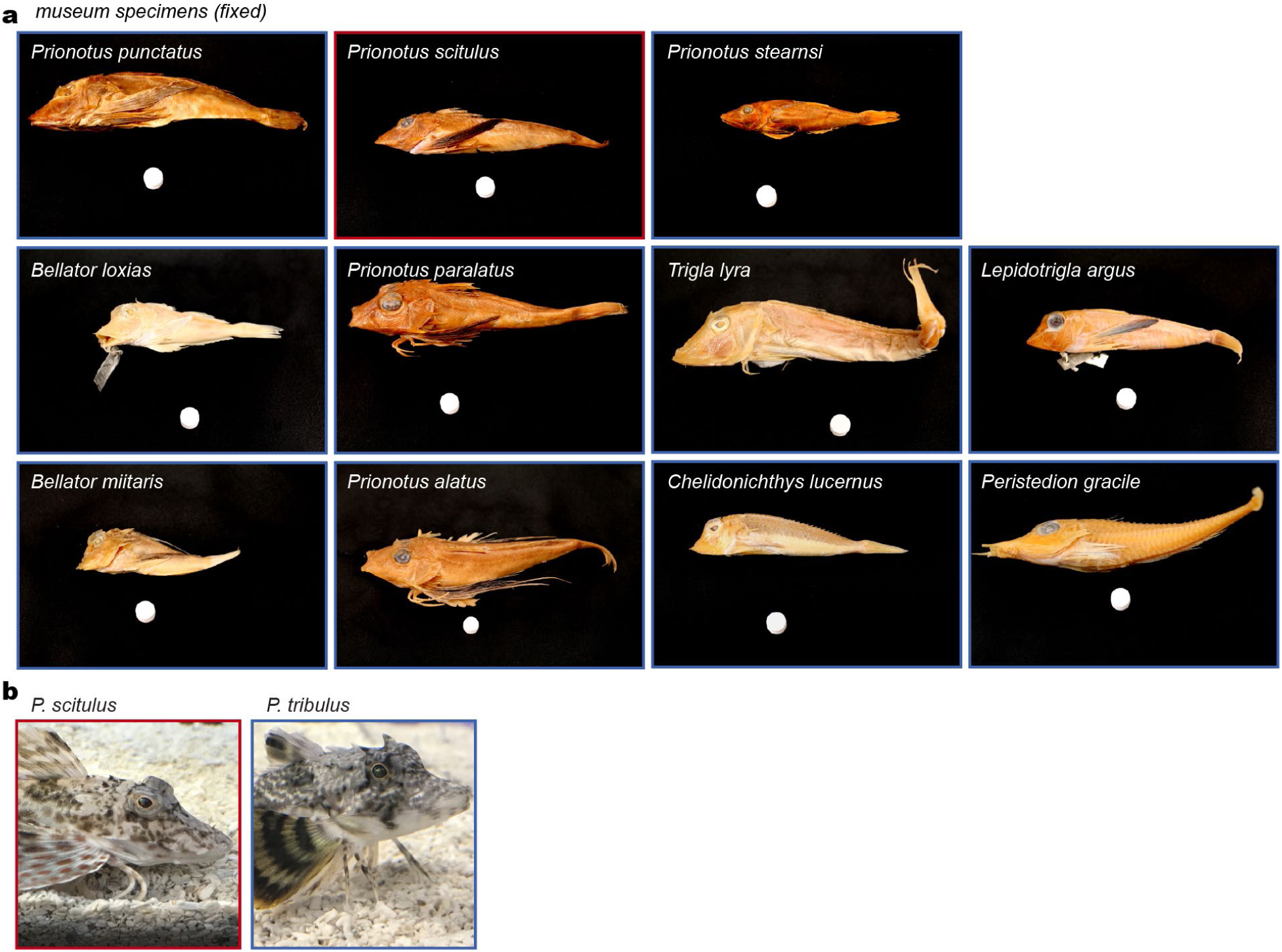
Comparative morphology. **a**, Museum specimens used for leg morphological analysis. Scale dot = 1 cm. **b**, Lateral views of the wild-caught *P. scitulus*, which had leg papillae, and *P. tribulus*, which lacked papillae, used in physiological and behavioral experiments. Red outline = papillae, blue outline = no papillae.

**Supplementary Movie 1.** Example of a digging sea robin (*P. carolinus*) “walking” around the tank and probing a mussel shell. Movie in real time.

## References

1. Russell, M., Grace, M. & Gutherz, E. Field guide to the searobins (Prionotus and Bellator) in the Western North Atlantic. (1992).

2. Bardach, J. E. & Case, J. Sensory capabilities of the modified fins of squirrel hake (*Urophycis chuss*) and Searobins (*Prionotus carolinus* and *P. evolans*). Copeia 1965, 194–206 (1965).

3. Manderson, J. P., Phelan, B. A., Bejda, A. J., Stehlik, L. L. & Stoner, A. W. Predation by striped searobin (*Prionotus evolans*, Triglidae) on young-of-the-year winter flounder (Pseudopleuronectes americanus, Walbaum): examining prey size selection and prey choice using field observations and laboratory experiments. J. Exp. Mar. Biol. Ecol. 242, 211–231 (1999).

4. Ross, S. T. Patterns of resource partitioning in searobins (Pisces: Triglidae). Copeia 1977, 561– 571 (1977).

5. Sazima, C. & Grossman, A. A non-digging zoobenthivorous fish attracts two opportunistic predatory fish associates. Neotropical Ichthyology 3, 445–448 (2005).

6. Silver, W. L. & Finger, T. E. Electrophysiological examination of a non-olfactory, non-gustatory chemosense in the searobin, *Prionotus carolinus*. J. Comp. Physiol. 154, 167–174 (1984).

7. Oike, H. et al. Characterization of ligands for fish taste receptors. J. Neurosci. 27, 5584–5592 (2007).

8. Schneider, E. R., Gracheva, E. O. & Bagriantsev, S. N. Evolutionary specialization of tactile perception in vertebrates. Physiology (Bethesda) 31, 193–200 (2016).

9. Saunders, C. J. & Silver, W. L. Anatomy and physiology of chemesthesis. in Chemesthesis 77–91 (John Wiley & Sons, Ltd, 2016). doi:10.1002/9781118951620.ch5.

10. Finger, T. E. Ascending spinal systems in the fish, Prionotus carolinus. Journal of Comparative Neurology 422, 106–122 (2000).

11. Hill, R. Z., Loud, M. C., Dubin, A. E., Peet, B. & Patapoutian, A. Piezo1 transduces mechanical itch in mice. Nature 607, 104–110 (2022).

12. Shin, S. M. et al. Peripheral sensory neurons and non-neuronal cells express functional Piezo1 channels. Mol Pain 19, 17448069231174315 (2023).

13. Li, X. et al. Human receptors for sweet and umami taste. Proceedings of the National Academy of Sciences 99, 4692–4696 (2002).

14. Nelson, G. et al. Mammalian sweet taste receptors. Cell 106, 381–390 (2001).

15. Morais, S. The physiology of taste in fish: potential implications for feeding stimulation and gut chemical sensing. Reviews in Fisheries Science & Aquaculture 25, 133–149 (2017).

16. Toda, Y., Okada, S. & Misaka, T. Establishment of a new cell-based assay to measure the activity of sweeteners in fluorescent food extracts. J. Agric. Food Chem. 59, 12131–12138 (2011).

17. Toda, Y. et al. Two distinct determinants of ligand specificity in T1r1/T1r3 (the umami taste receptor). J Biol Chem 288, 36863–36877 (2013).

18. Yuschak, P. Fecundity, eggs, larvae and osteological development of the striped searobin, (*Prionotus evolans*) (Pisces, Triglidae). J. Northw. Atl. Fish. Sci. 6, 65–85 (1985).

19. Yuschak, P. & Lund, W. A. Eggs, larvae and osteological development of the Northern Searobin, *Prionotus carolinus* (Pisces, Triglidae). J. Northw. Atl. Fish. Sci. 5, 1–15 (1984).

20. Morrill, A. D. The pectoral appendages of *Prionotus* and their innervation. Journal of Morphology 11, 177–192 (1895).

21. Gerhardt, Greg. & Adams, R. N. Determination of diffusion coefficients by flow injection analysis. Anal. Chem. 54, 2618–2620 (1982).

22. Casalini, T., Salvalaglio, M., Perale, G., Masi, M. & Cavallotti, C. Diffusion and aggregation of sodium fluorescein in aqueous solutions. J. Phys. Chem. B 115, 12896–12904 (2011).

23. Ma, Y., Zhu, C., Ma, P. & Yu, K. T. Studies on the diffusion coefficients of amino acids in aqueous solutions. J. Chem. Eng. Data 50, 1192–1196 (2005).

24. Crank, J. The mathematics of diffusion. (Clarendon Press, 1975).

25. Krueger, F. et al. FelixKrueger/TrimGalore: v0.6.10. (2023) doi:10.5281/zenodo.7598955.

26. Grabherr, M. G., et al. Trinity: reconstructing a full-length transcriptome without a genome from RNA-Seq data. Nat Biotechnol 29, 644–652 (2011).

27. Bray, N. L., Pimentel, H., Melsted, P. & Pachter, L. Near-optimal probabilistic RNA-seq quantification. Nat Biotechnol 34, 525–527 (2016).

28. Buchfink, B., Reuter, K. & Drost, H.-G. Sensitive protein alignments at tree-of-life scale using DIAMOND. Nat Methods 18, 366–368 (2021).

29. Meade, M. E., Roginsky, J. E. & Schulz, J. R. Primary cell culture of adult zebrafish spinal neurons for electrophysiological studies. J. Neurosci. Methods 322, 50–57 (2019).

30. Portnoy, D. S. et al. Molecular phylogenetics of New World searobins (Triglidae; Prionotinae). Molecular Phylogenetics and Evolution 107, 382–387 (2017).

